# Enhanced negative emotional processing limits behavioral flexibility in a mouse model of autism spectrum disorder

**DOI:** 10.1101/2021.05.31.446503

**Authors:** Miru Yun, Eunjoon Kim, Min Whan Jung

**Author notes:** Correspondence to: EK and MWJ.

## Abstract

Impaired behavioral flexibility might underlie some of the symptoms associated with autism spectrum disorder (ASD). We investigated whether and how behavioral flexibility is impaired in a mouse model of ASD by testing *Shank2*-knockout (*Shank2*-KO) mice in reversal learning. *Shank2*-KO mice were trained in probabilistic classical conditioning with two odor cues paired with water and air puff. Upon the reversal of cue-outcome contingency, *Shank2*-KO mice were significantly slower than wild-type mice in reversing their anticipatory licking responses. *Shank2*-KO mice also showed stronger anticipatory eye closure responses than wild-type mice to the air puff, raising a possibility that the impairment might be because of enhanced negative emotional processing. Indeed, *Shank2-*KO mice showed intact reversal learning when the strong air puff was replaced with a mild air puff. *Shank2-*KO mice also showed intact reversal learning between two odor cues predicting rewards with different probabilities. These results indicate that enhanced negative emotional processing suppresses reversal learning despite of intact capability to learn cue-outcome contingency changes in *Shank2*-KO mice in our behavioral settings. Our findings suggest that behavioral flexibility may be seriously limited by abnormal emotional processing in ASD.

## Introduction

Autism spectrum disorders (ASD) are developmental disorders that are associated with a diverse array of symptoms including impaired social interaction and communication as well as repetitive and restrictive patterns of behavior (American Psychiatric Association, 2013). Many genes implicated in ASD are expressed broadly in the cerebral cortex and, therefore, their mutations can potentially lead to aberrant circuit development in widespread cortical areas (Hashem et al., 2020; Mundy, 2003; Yan & Rein, 2021). Given that the cerebral cortex plays a key role in adaptive control of behavior, mutations in genes playing important roles in cortical development may well lead to compromised behavioral flexibility in adults. In this respect, ASD patients often show impairments in the tasks requiring cognitive flexibility such as the Wisconsin card sorting test (Hill, 2004; Leung & Zakzanis, 2014), and impaired cognitive flexibility has been proposed to underlie repetitive and restrictive patterns of behavior, a core symptom of ASD (Geurts, Corbett, & Solomon, 2009). ASD patients also show impairments in reversal learning with their performance correlated with clinical ratings of restricted and repetitive behaviors (D’Cruz et al., 2013) or everyday symptoms of behavioral inflexibility (South, Newton, & Chamberlain, 2012). In addition, impaired behavioral flexibility has been reported in mouse models of ASD. Two mouse models of ASD, BTBR T+ tf/J (BTBR) and C58 mice, showed impaired spatial reversal learning with their performance negatively correlated with repetitive behavior (D.A. Amodeo, Jones, Sweeney, & Ragozzino, 2012; Whitehouse, Curry-Pochy, Shafer, Rudy, & Lewis, 2017). Collectively, albeit limited, studies have found impaired behavioral flexibility and its correlations with repetitive and restrictive behaviors in ASD patients and animal models, raising a possibility that repetitive and restrictive patterns of behavior associated with ASD might be manifestations of impaired flexibility.

Shank2 (SH3 and multiple ankyrin repeat domains 2), which is a multi-domain scaffolding protein enriched in the postsynaptic density of excitatory synapses (Sheng & Kim, 2000, 2011), is strongly implicated in ASD (Abrahams et al., 2013). Genetic variations of *SHANK2* gene have been identified in ASD (Aspromonte et al., 2019; Bai et al., 2018; Berkel et al., 2010; Berkel et al., 2012; Chilian et al., 2013; Costas, 2015; Guilmatre, Huguet, Delorme, & Bourgeron, 2014; Homann et al., 2016; Leblond et al., 2012; Leblond et al., 2014; Liu et al., 2013; Lu, Mu, Osborne, & Cordner, 2018; Mossa, Giona, Pagano, Sala, & Verpelli, 2018; S Peykov et al., 2015; S. Peykov et al., 2015; Pinto et al., 2010; Prasad et al., 2012; Rauch et al., 2012; Sanders et al., 2012; Satterstrom et al., 2020; Schluth-Bolard et al., 2013; Wang et al., 2020; Wischmeijer et al., 2010; Yuen et al., 2017; Zhou et al., 2019). Moreover, mutations/deletions in *Shank2* gene lead to a diverse array of behavioral changes in mice, and some of them show impaired social interaction and repetitive behavior, two core symptoms of ASD (Eltokhi, Rappold, & Sprengel, 2018; Ha et al., 2016; Kim et al., 2018; Lim et al., 2017; Pappas et al., 2017; Peter et al., 2016; Schmeisser et al., 2012; Won et al., 2012). The mice with alterations in *Shank2* gene have been used widely to investigate neurobiological mechanisms of ASD and, as a result, substantial amounts of behavioral and neural data have been accumulated. However, the relationship between Shank2 mutations and behavioral flexibility is largely unknown. In the present study, we investigated whether and how behavioral flexibility is compromised in *Shank2* homozygous knock-out (*Shank2*-KO) mice. For this, we subjected *Shank2-*KO mice to reversal learning using a probabilistic classical conditioning paradigm. The results indicate that *Shank2-*KO mice have intact capability to update changes in cue-outcome contingency, but display abnormally heightened negative emotional responses that limit behavioral flexibility under certain circumstances.

## Results

We used male *Shank2-*KO mice with deletions in exons 6 and 7 of *Shank2* gene which mimic the microdeletion of exons 6 and 7 in human *SHANK2* gene identified in ASD (Won et al., 2012). The mice were trained in a probabilistic classical conditioning task under head fixation (**Figure 1a**) as previously described (Jeong, Kim, Song, Paik, & Jung, 2020). Two different odor cues (randomly chosen in a given trial) were paired with an appetitive outcome (water) or an aversive outcome (air puff) with given probabilities (**Figure 1b**).

**Figure 1.**
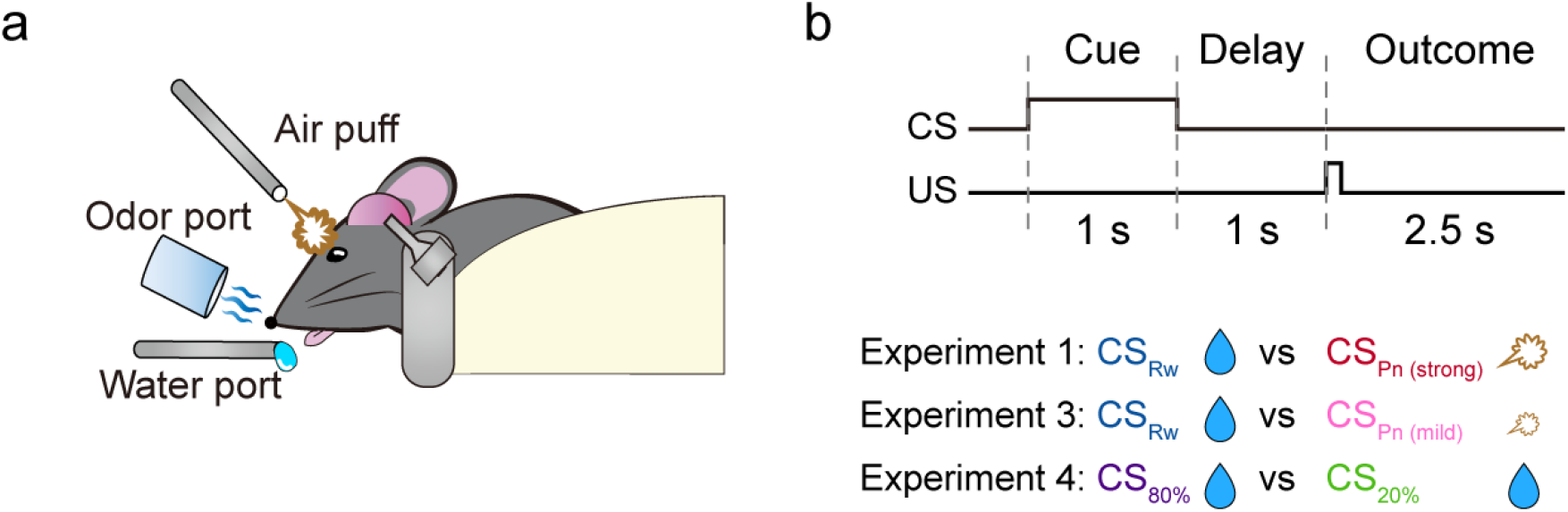
Behavioral task. (**a**) The experimental setting. Head-fixed mice performed a probabilistic classical conditioning task. (**b**) Task schematic. A conditioned stimulus (CS; odor) was delivered for 1 s and, after a 1-s delay, an unconditioned stimulus (US), either appetitive (water, 6 μl) or aversive (air puff), was delivered in a probabilistic manner. Each CS was paired with 75% water (CS_Rw_) or 75% strong air puff (CS_Pn(strong)_) in Experiment 1, 75% water or 75% mild air puff (CS_Pn(mild)_) in Experiment 3, and 80% water (CS_80%_) or 20% water (CS_20%_) in Experiment 4.

### Experiment 1: Appetitive/aversive reversal learning

In Experiment 1, one odor cue was paired with a small amount of water (6 μl) and another with an air puff (100 ms, 3 psi) each with 75% probability (**Figure 2a**). Wild-type (WT; n = 10) and *Shank2-*KO mice (n = 10) were trained in this probabilistic appetitive/aversive conditioning task for three daily sessions (acquisition phase; 400 trials per session). We used anticipatory lick rate during the delay period (1 s) as an index for discrimination between reward-predicting and punishment-predicting cues throughout the study. The anticipatory lick rate in CS_Rw_ trials increased during training. Two-way mixed ANOVA indicated a significant main effect of session (*F_(2, 36)_* = 4.629, *p* = 0.016) without a significant main effect of animal group (*F_(1, 18)_* = 2.695, *p* = 0.118) or session×animal group interaction (*F_(2, 36)_* = 0.347, *p* = 0.709; **Figure 2b**). In contrast, the anticipatory lick rate in CS_Pn_ trials did not change significantly during training (main effect of session, *F_(2, 36)_* = 2.0274, *p* = 0.146; main effect of animal group, *F_(1, 18)_* = 1.311, *p* = 0.267; session×animal group interaction, *F_(2, 36)_* = 1.088, *p* = 0.348; **Figure 2c**). During the last acquisition session, as shown by two sample sessions in **Figure 2a**, the anticipatory lick rate during the delay period was significantly higher in CS_Rw_ than CS_Pn_ trials in both animal groups (WT mice, 2.7±0.5 and 0.6±0.1 Hz, respectively; paired *t*-test, *t_(9)_* = 3.804: *p* = 0.001; *Shank2-*KO mice, 3.9±0.6 and 1.0±0.2 Hz, respectively; *t_(9)_* = 4.563: *p* = 2.4×10^−4^). These results indicate intact motivation and capability to learn the cue-outcome contingency in *Shank2-* KO mice.

**Figure 2.**
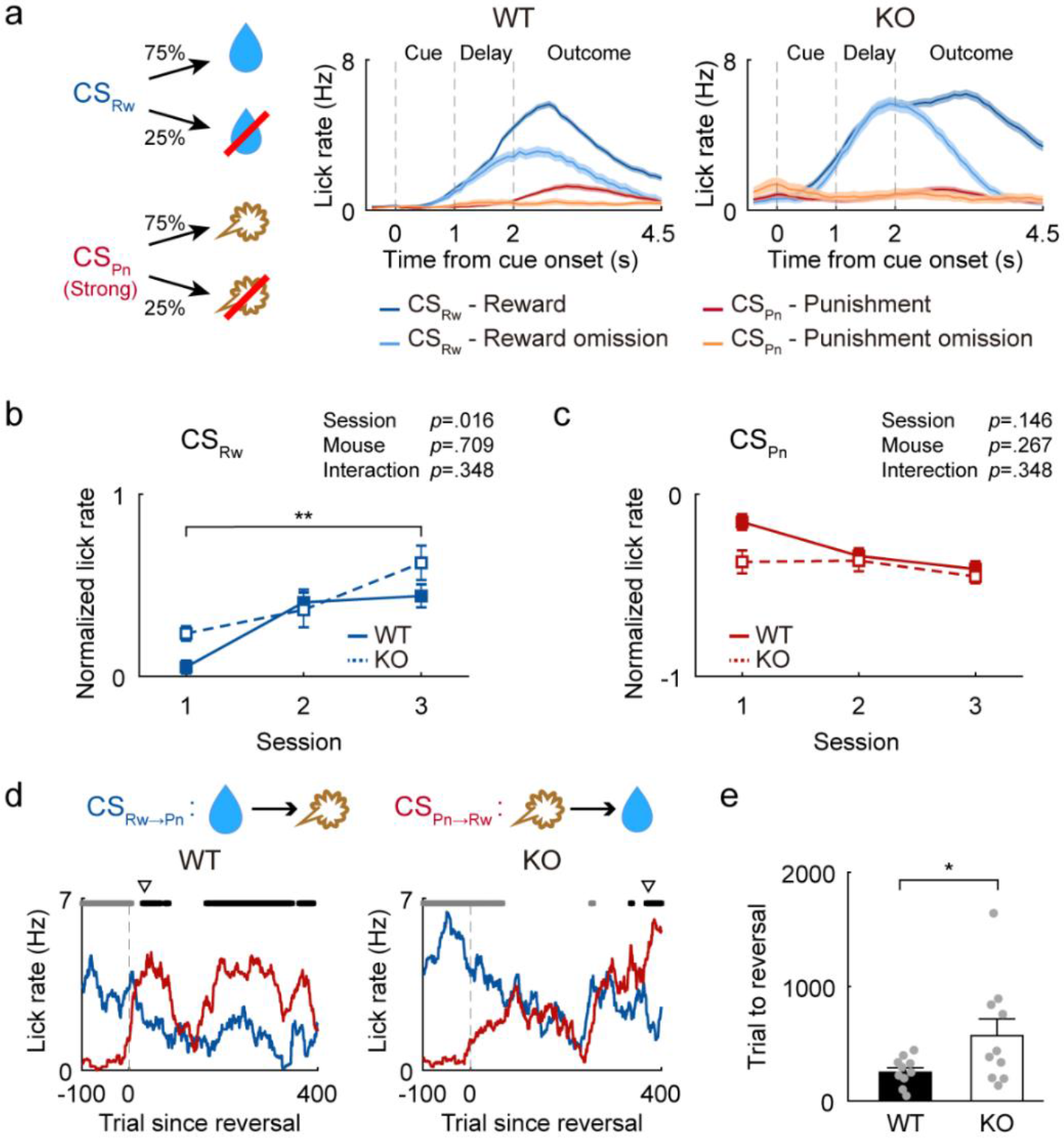
Impaired reversal learning in *Shank2-*KO mice. Results of Experiment 1. (**a**) Left, two odor cues, CS_Rw_ and CS_Pn_, were paired with 75% reward (water, 6 μl; blue) and 75% strong air puff (100 ms, 3 psi; red), respectively. Right, samples for licking responses (lick density functions, σ = 100 ms) of WT (left) and *Shank2-*KO (right) mice during the last acquisition session. Trials were grouped according to CS and outcome. Shading, SEM across trials. (**b** and **c**) Z-normalized anticipatory lick rates (delay-period lick rates) of WT (solid line) and *Shank2-*KO (dashed line) mice in CS_Rw_ (**b**) and CS_Pn_ (**c**) trials during initial acquisition. *P*-values, outcomes of two-way mixed ANOVA. \*\**p* < 0.01, post-hoc Bonferroni test. (**d**) Sample reversal-learning sessions. Blue and red lines indicate anticipatory licking responses to CS_Rw→Pn_ and CS_Pn→Rw_ cues, respectively, in moving average of 25 trials. Gray and black squares on top, significantly higher anticipatory licking response to CS_Rw→Pn_ (gray) or CS_Pn→Rw_ (black) cue than the other in a moving window of 25 trials (*p* < 0.05, paired *t*-test). Triangles on top indicate the first trial since cue-outcome contingency reversal that exceeded the reversal criterion. (**e**) The number of trials to exceed the reversal criterion. Gray circles, individual animal data; bar graphs and error bars, mean and SEM. \**p* < 0.05, *t*-test.

The mice were then subjected to reversal learning. We reversed cue-outcome contingency so that the previous reward-predicting cue becomes the punishment-predicting cue after reversal (CS_Rw→Pn_) and vice versa (CS_Pn→Rw_). All mice were trained until they reach the reversal criterion over 1~5 sessions (400 trials per daily session). The number of trials to reach the reversal criterion differed significantly between WT and *Shank2-*KO mice (250.0±39.7 and 570.7±146.6 trials, respectively; *t*-test, *t_(18)_* = 2.112, *p* = 0.049; **Figure 2d,e**). The mice sometimes consumed water in the subsequent trial rather than during the inter-trial interval following a rewarded trial. To rule out the influence of such invalid anticipatory licks, we determined the number of trials to reversal criterion after deleting the trials during which the mice consumed water delivered in the previous trial (226 out of 8213 trials; 2.75%). This analysis also yielded a significant difference in the number of trials to reach the reversal criterion between WT and *Shank2-*KO mice (250.1±39.7 and 571.2±146.7 trials, respectively; *t_(18)_* = 2.112, *p* = 0.049). These results indicate slower reversal learning in *Shank2-*KO than WT mice.

### Experiment 2: Eye closure responses to air puff

ASD is associated with atypical sensory responses and heightened anxiety (Chen et al., 2020; Kazdoba et al., 2015; Vasa et al., 2014). We therefore examined whether WT and *Shank2-*KO mice show differential eye closure responses to the air puff using separate groups of WT (n = 8) and *Shank2-*KO (n = 8) mice. Specifically, we delivered the identical air puff used in Experiment 1 (100 ms, 3 psi) without any preceding sensory cue (ITI, 9~11 s, uniform random distribution) and measured the fraction of eye closure before, during, and after air puff delivery (**Figure 3a**). Eye closure response did not differ significantly between WT and *Shank2-*KO mice immediately after air puff delivery (1-s time window since air puff onset; *t*-test, *t_(14)_* = 1.484, *p* = 0.160; **Figure 3b**). However, eye closure response differed significantly between WT and *Shank2-*KO mice before (1.5-s time window before air puff onset; *t_(14)_* = 2.380, *p* = 0.032) and after (between 2.5 and 4 s since air puff onset; *t_(14)_* = 2.157, *p* = 0.049) air puff delivery (**Figure 3b**). These results indicate differential anticipatory eye closure responses between WT and *Shank2-*KO mice.

**Figure 3.**
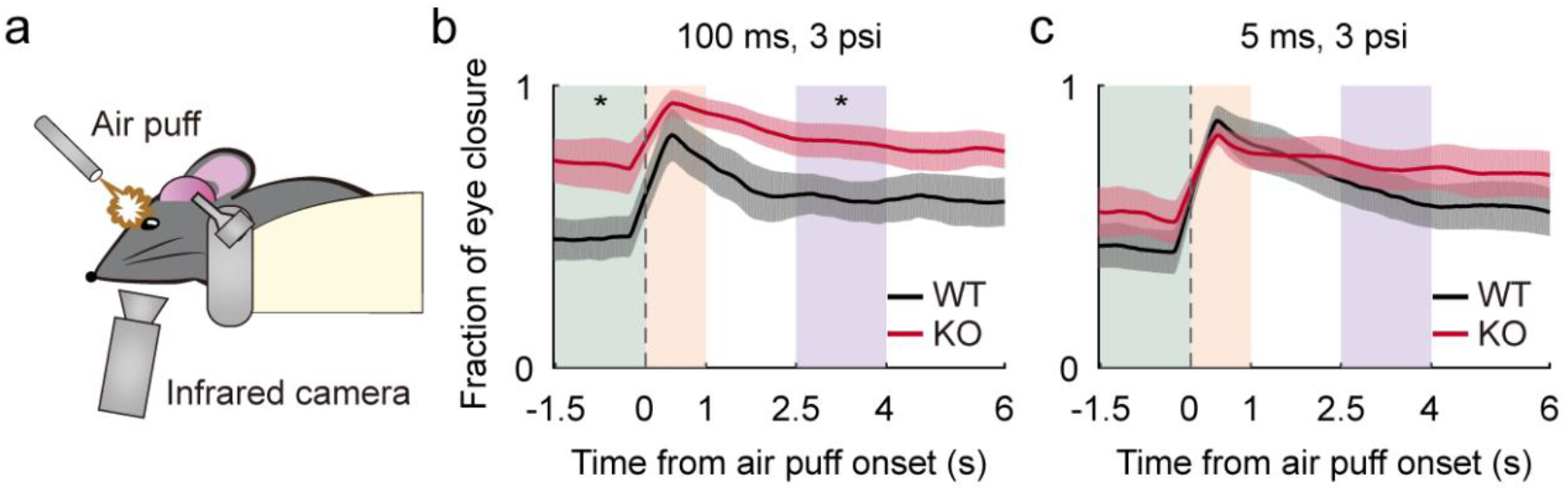
Eyelid closure response to air puff differs between WT and *Shank2-*KO mice. Results of Experiment 2. (**a**) Eyelid closure responses to air puff were estimated by measuring pupil diameter with an infrared camera. (**b**) Eyelid closure responses (average of 5 trials) of WT (black) and *Shank2-*KO (red) mice (shading, SEM across mice) to the strong (100 ms, 3 psi) air puff used in Experiment 1. Shaded rectangles indicate time periods before (green, 1.5 s before air puff onset), immediately after (orange, 1 s after air puff onset), and after air puff delivery (purple, between 2.5 and 4 s after air puff onset). \**p* < 0.05, *t*-test. (**c**) Eyelid closure responses to the mild (5 ms, 3 psi) air puff used in Experiment 3.

The above results raise a possibility that abnormal emotional responses to air puff may negatively affect reversal learning in *Shank2-*KO mice. We therefore examined the relationship between air puff strength and eye closure response by systematically varying the duration and intensity of air puff (15 combinations except for the original one (100 ms, 3 psi)). Eye closure responses immediately after air puff delivery (1 s) did not differ significantly between the two animal groups in all combinations of air puff duration and intensity (*t*-test, *p*-values > 0.05; **Figure 3-supplementary figure 1**). However, anticipatory eye closure responses before (1.5-s time window before air puff onset) and after air puff delivery (between 2.5 and 4 s since air puff onset) differed significantly between the two animal groups in some intermediate-strength combinations. Mild air puffs induced similarly low levels of anticipatory eye closure responses (**Figure 3c**), and powerful air puffs induced similarly high levels of anticipatory eye closure responses (**Figure 3-supplementary figure 1a**) in WT and *Shank2-*KO mice. However, to some intermediate-strength air puffs (’strong’ air puffs), the anticipatory eye closure response was significantly stronger in *Shank2-*KO than WT mice (**Figure 3-supplementary figure 1** and **Figure 3-supplementary table 1**). These results suggest altered emotional processing in *Shank2-*KO mice. Based on these results, we chose to use the mildest air puff (5 ms, 3 psi; **Figure 3c**) to further examine reversal learning of *Shank2-*KO mice.

### Experiment 3: Appetitive/aversive reversal learning with a mild air puff

In Experiment 3, WT (n = 10) and *Shank2-*KO mice (n = 10) were trained in the appetitive/aversive conditioning in the same manner as in Experiment 1 except that a mild air puff (5 ms, 3 psi), to which the two animal groups show similar anticipatory eye closure responses, was used (**Figure 4a**). The anticipatory lick rate during the delay period increased during the initial training with no significant difference between the animal groups in CS_Rw_ trials (two-way mixed ANOVA, main effect of session, *F_(2, 36)_* = 3.313, *p* = 0.048; main effect of mouse group, *F_(1, 18)_* = 0.270, *p* = 0.070; mouse group×session interaction, *F_(2, 36)_* = 1.162, *p* = 0.324; **Figure 4b**). In CS_Pn_ trials, the anticipatory lick rate did not change significantly during training, and *Shank2-*KO mice showed overall higher anticipatory lick rates than WT mice (main effect of session, *F_(2, 36)_* = 1.863, *p* = 0.170; main effect of mouse group, *F_(1, 18)_* = 12.453, *p* = 0.002; mouse group×session interaction, *F_(2, 36)_* = 0.954, *p* = 0.395; **Figure 4c**). During the last acquisition session, as shown by two sample sessions in **Figure 4a**, the anticipatory lick rate during the delay period was significantly higher in CS_Rw_ than CS_Pn_ trials in both animal groups (WT mice, 3.3±0.4 and 0.4±0.1 Hz, respectively; paired *t*-test, *t_(9)_* = 6.469: *p* = 4.4×10^−6^; *Shank2-*KO mice, 3.3±0.6 and 1.2±0.3 Hz, respectively; *t_(9)_* = 3.359: *p* = 0.004). These results indicate intact learning capability of *Shank2-*KO mice for discriminating between reward-predicting and mild punishment-predicting cues.

**Figure 4.**
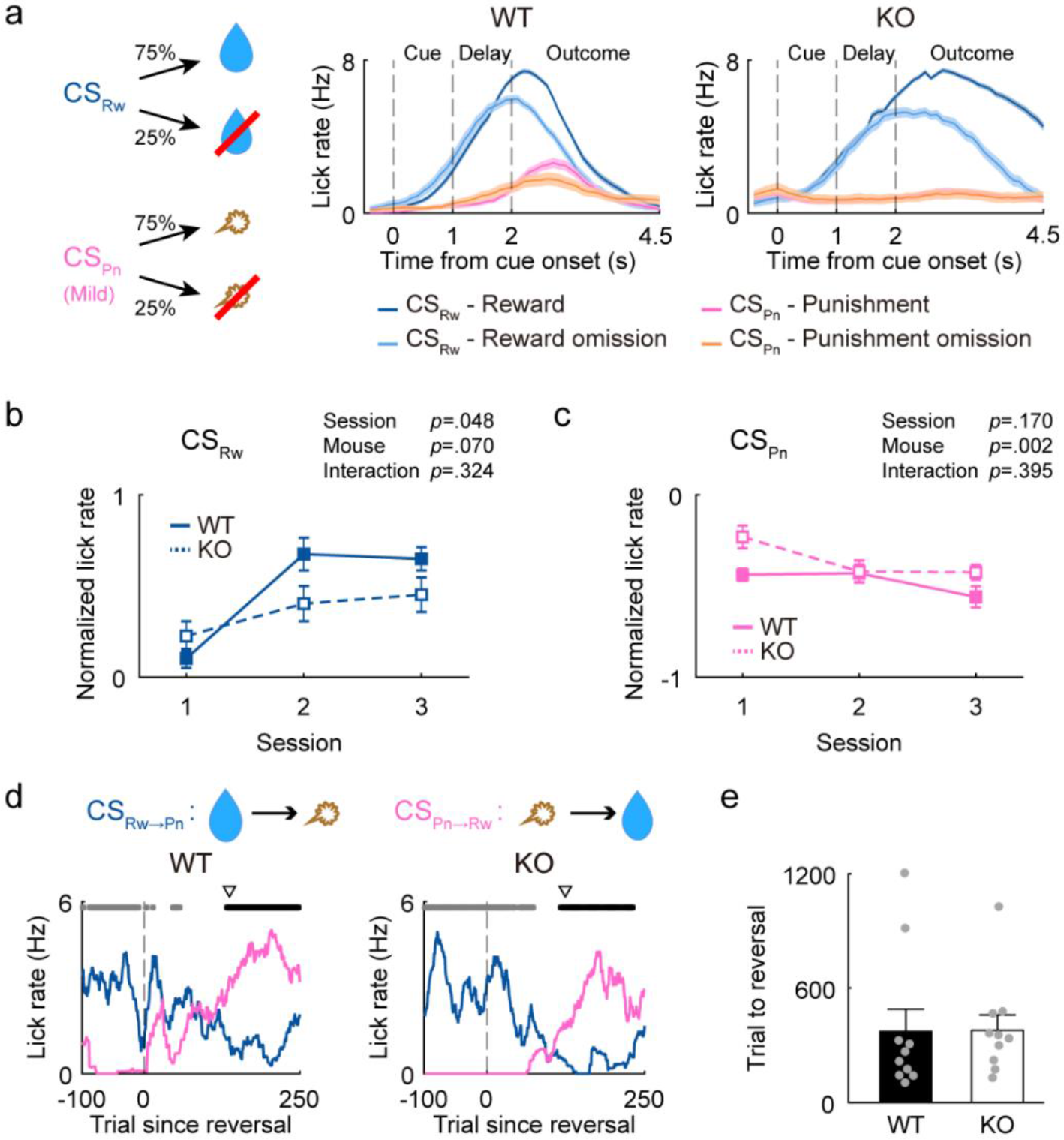
Intact reversal learning of *Shank2-*KO mice with the use of a mild air puff. Results of Experiment 3. (**a**) Left, Two odor cues, CS_Rw_ and CS_Pn_, were paired with 75 % reward (water, 6 μl; blue) and 75 % mild air puff (5 ms, 3 psi; pink), respectively. Right, Samples for licking responses during the last acquisition session. (**b** and **c**) Z-normalized anticipatory lick rates in CS_Rw_ (**b**) and CS_Pn_ (**c**) trials during initial acquisition. (**d**) Sample reversal-learning sessions. Blue and pink lines denote anticipatory licking responses to CS_Rw→Pn_ and CS_Rw→Pn_ cues, respectively. (**e**) The number of trials to exceed the reversal criterion. The same format as in **Figure 2**.

We then tested the mice in reversal learning. The test procedure was identical to that in Experiment 1. Unlike in Experiment 1, the number of trials to reach the reversal criterion did not differ significantly between WT and *Shank2-*KO mice (373.6±117.5 and 379.0±79.9 trials, respectively; *t*-test, *t_(18)_* = 0.038, *p* = 0.970; **Figure 4d,e**). Similar results were obtained when we performed the same analysis after removing the trials during which mice consumed water delivered in the previous trial (114 out of 7697 trials; 1.48%; 390.7±113.9 and 379.0±79.9 trials to reversal criterion in WT and *Shank2-*KO mice, respectively; *t_(18)_* = 0.084, *p* = 0.934). These results show intact reversal learning of *Shank2-*KO mice, suggesting that impaired reversal learning in Experiment 1 might be because of altered emotional responses of *Shank2-*KO mice to a strong air puff.

### Experiment 4: Appetitive/appetitive reversal learning

To further confirm intact behavioral flexibility of *Shank2-*KO mice in the absence of strong negative emotions, we tested WT (n = 10) and *Shank2-*KO (n = 10) mice in reversal learning using only appetitive outcomes. In Experiment 4, two different odor cues were paired with the same amount of water (6 μl), but with two different probabilities (80 and 20%; CS_80%_ and CS_20%_; **Figure 5a**). The anticipatory lick rate during the delay period increased over three training sessions in CS_80%_ trials (two-way mixed ANOVA, main effect of session, *F_(2, 36)_* = 6.304, *p* = 0.005; main effect of mouse group, *F_(1, 18)_* = 3.266, *p* = 0.088; mouse group×session interaction, *F_(2, 36)_* = 1.277, *p* = 0.291; **Figure 5b**), but not in CS_20%_ trials (main effect of session, *F_(2, 36)_* = 1.940, *p* = 0.158; main effect of mouse group, *F_(1, 18)_* = 2.246, *p* = 0.151; mouse group×session interaction, *F_(2, 36)_* = 0.052, *p* = 0.949; **Figure 5c**). During the last acquisition session, as shown by two sample sessions in **Figure 5a**, the anticipatory lick rate was significantly higher in CS_80%_ than CS_20%_ trials in both animal groups (WT mice, 3.1±0.3 and 1.6±0.2 Hz, respectively; paired *t*-test, *t_(9)_* = 3.991, *p* = 8.6×10^−4^; *Shank2-*KO mice, 3.7±0.6 and 2.1±0.2 Hz, respectively; *t_(9)_* = 2.477, *p* = 0.023). These results indicate that both animal groups successfully learned the cue-outcome contingency.

**Figure 5.**
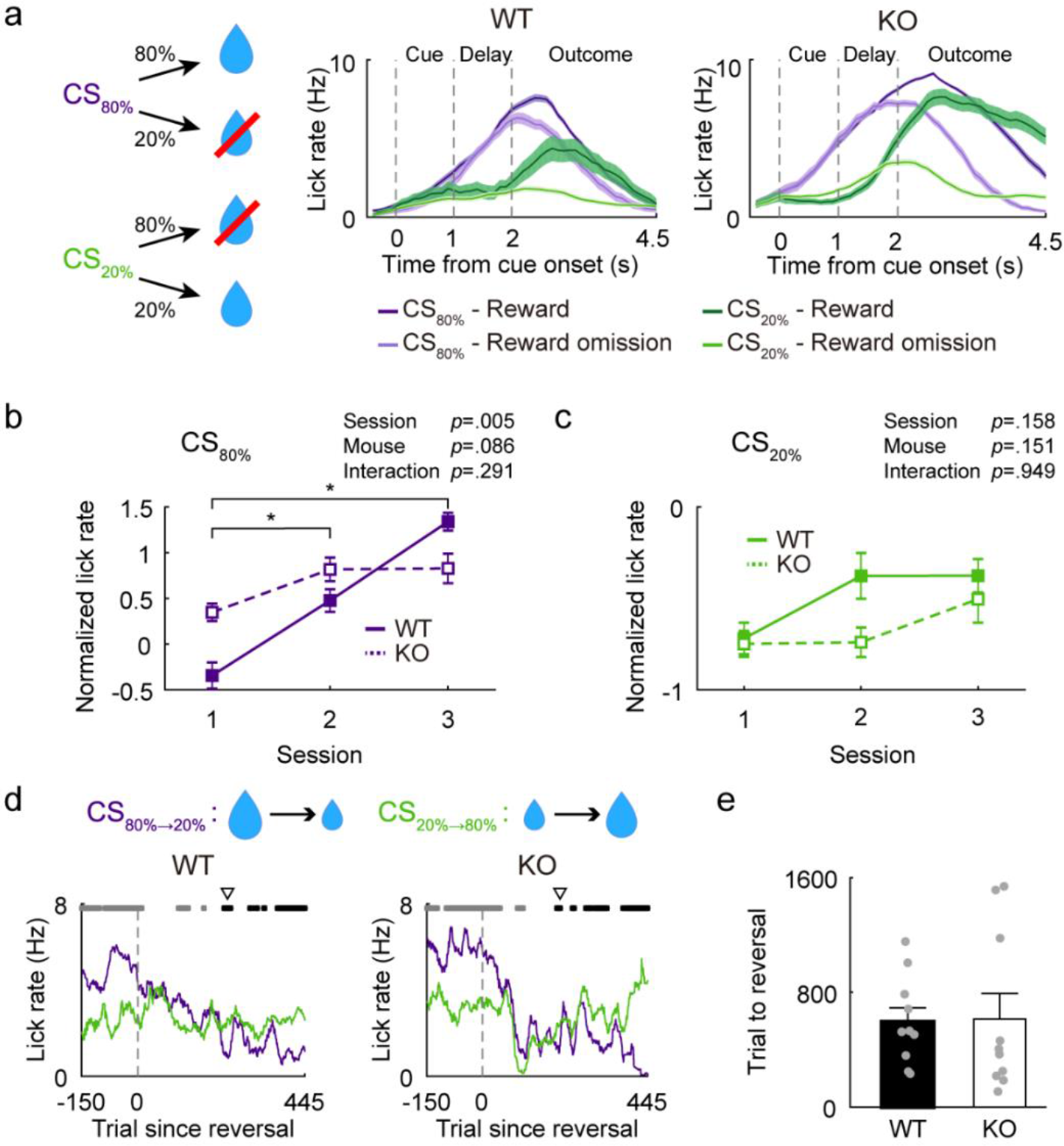
Intact learning of *Shank2-*KO mice in reward probability reversal. Results of Experiment 4. (**a**) Left, Two odor cues, CS_80%_ (purple) and CS_20%_ (green), were paired with 80% and 20% reward (water, 6 μl), respectively. Right, Sample licking responses during the last acquisition session. (**b** and **c**) Z-normalized anticipatory lick rates in CS_80%_ (**b**) and CS_20%_ (**c**) trials during initial acquisition. \**p* < 0.05, post-hoc Bonferroni test. (**d**) Sample reversal-learning sessions. Purple and green lines denote anticipatory licking responses to CS_80%→20%_ and CS_20%→80%_ cues, respectively. (**e**) The number of trials to exceed the reversal criterion. The same format as in **Figure 2**.

The mice were then tested in reversal learning in the same manner as in Experiments 1 and 3 except that two appetitive outcomes were used (**Figure 5d**). The number of trials to reach the reversal threshold did not differ significantly between the two animal groups (620.1±97.7 and 633.6±189.2 trials, respectively; *t*-test, *t_(18)_* = 0.063, *p* = 0.950; **Figure 5e**). Similar results were obtained when we repeated the analysis after deleting the trials during which mice consumed water delivered in the previous trial (287 out of 12537 trials; 2.29%; 604.9±96.4 and 463.7±113.0 trials to reversal criterion in WT and *Shank2-*KO mice, respectively; *t_(18)_* = 0.950, *p* = 0.355). These results further show intact reversal learning of *Shank2-*KO mice in the absence of strong aversive outcomes.

## Discussion

We examined behavioral flexibility of *Shank2-*KO mice by testing their capability for reversal learning in a probabilistic classical conditioning task using anticipatory lick rate as an index for learning. Compared to WT mice, *Shank2-*KO mice showed significantly slower reversal learning when a water reward and a strong air puff were used as appetitive and aversive outcomes, respectively (Experiment 1). However, *Shank2-*KO mice showed stronger eye closure responses than WT mice to the anticipated air puff (Experiment 2), raising a possibility that the impaired reversal learning in Experiment 1 might be because of altered emotional processing. Indeed, when we replaced the strong air puff with a mild air puff that induced similar eye close responses between *Shank2-*KO and WT mice, *Shank2-*KO mice showed intact reversal learning (Experiment 3). Furthermore, *Shank2-*KO mice showed intact reversal learning between two cues predicting rewards with different probabilities (Experiment 4). Together, these results are consistent with enhanced negative emotional processing, but intact learning of cue-outcome contingency changes in *Shank2-*KO mice in our behavioral settings. A previous study has found delayed reversal learning in ASD patients in a fear conditioning paradigm in which an air puff was used as an unconditioned stimulus (South et al., 2012). Our results raise a possibility that ASD patients may show intact reversal learning if the unconditioned stimulus is replaced with a mild air puff, which remains to be tested. Numerous studies indicate the role of the orbitofrontal cortex in reversal learning (L. Amodeo, McMurray, & Roitman, 2017; Ragozzino, 2007). We also have shown that the intact medial prefrontal cortex is needed for probabilistic reversal learning in mice (Jeong et al., 2020). Our results suggest that the neural machinery needed to keep track of changes in cue-outcome contingency as well as that needed to control behavior according to such evaluations are likely to be intact in these brain structures of *Shank2-* KO mice.

On the one hand, eye closure responses were similar between *Shank2-*KO and WT mice immediately after an air puff was delivered. This suggests that abnormal sensory processing, which is often observed in ASD patients (Balasco, Provenzano, & Bozzi, 2019; Marco, Hinkley, Hill, & Nagarajan, 2011) and ASD animal models (Balasco et al., 2019; Orefice et al., 2016), or abnormal reactivity to an aversive stimulus is unlikely to account for the reversal learning deficit of *Shank2-*KO mice observed in Experiment 1. On the other hand, *Shank2-*KO mice showed significantly stronger eye closure responses than WT mice before and after air puff delivery. Fear and anxiety are the emotional states that evoke a defense state against a threat (Davis, Walker, Miles, & Grillon, 2010; Tovote, Fadok, & Lüthi, 2015). Fear refers to an emotional response to a real or perceived imminent threat, whereas anxiety is often elicited by a less specific and an expected (not real) threat (American Psychiatric Association, 2013; Davis et al., 2010). Therefore, our results suggest that enhanced anxiety, rather than enhanced fear, may underlie slower reversal learning of *Shank2-*KO mice in Experiment 1. About 40% of children with ASD are comorbid with anxiety disorders, and the level of their anxiety is positively correlated with the degree of repetitive behavior (Evans, Canavera, Kleinpeter, Maccubbin, & Taga, 2005; Jitlina et al., 2017; Rodgers, Glod, Connolly, & McConachie, 2012; Simonoff et al., 2008; South et al., 2012; Top Jr et al., 2016; Vasa et al., 2014; Wigham, Rodgers, South, McConachie, & Freeston, 2015). Also, numerous ASD model mice, including *Shank2-*KO mice, show elevated anxiety (Kazdoba et al., 2015; Silverman, Yang, Lord, & Crawley, 2010; Won et al., 2012). Previous studies have found impaired behavioral flexibility in human patients with anxiety disorder (Ansari, Derakshan, & Richards, 2008; Eysenck, Derakshan, Santos, & Calvo, 2007; Lyche, Jonassen, Stiles, Ulleberg, & Landro, 2010) and mice with stress-induced enhanced anxiety (Bondi, Rodriguez, Gould, Frazer, & Morilak, 2008; George et al., 2015; Park & Moghaddam, 2017). Our results, together with these findings, raise the possibility that enhanced anxiety may limit behavioral flexibility and thereby enhance repetitive and restrictive behaviors in ASD.

Our findings are at odds with the previous studies that showed impaired spatial reversal learning in ASD patients (D’Cruz et al., 2013) and ASD model mice (D. A. Amodeo et al., 2012; Whitehouse et al., 2017) using only appetitive outcomes. One possible explanation would be that behavioral flexibility is differently compromised across ASD subjects (*Shank2-*KO mice versus BTBR and C58 mice and ASD patients). For example, *Shank2-* KO mice, but not BTBR and C58 mice and ASD patients, may have intact capability to update changes in cue-outcome contingency. Alternatively, the different results may be because of different experimental procedures rather than differences in ASD subjects. We tested head-fixed mice in a classical conditioning paradigm, whereas ASD patients and BTBR and C58 mice were freely moving and tested in an instrumental conditioning paradigm (D. A. Amodeo et al., 2012; Whitehouse et al., 2017). Hence, unlike in our study, the subjects were allowed to make a choice freely in these studies. Reward was delivered probabilistically in these studies and, notably, BTBR and C58 mice showed intact reversal learning when reward was delivered in an all-or-none manner. Anxiety is elevated under uncertainty (Grupe & Nitschke, 2013; Hirsh, Mar, & Peterson, 2012). Also, anxiety disorder patients prefer to play a passive rather than an active role in decision making (Anderson, Arnold, Angus, & Bryce, 2009). These results raise the possibility that uncertain outcomes under a free-choice condition may have elevated anxiety so as to impair reversal learning of ASD patients and BTBR and C58 mice in these studies. Note that conflicting results have been reported regarding impaired reversal learning in young children (preschoolers) with ASD with certain reward delivery (Coldren & Halloran, 2003; Griffith, Pennington, Wehner, & Rogers, 1999; McEvoy, Rogers, & Pennington, 1993). Given that perseveration was the main source of errors, impaired reversal in some of these studies may be because of delayed development of executive functions, such as inhibitory control, rather than enhanced anxiety. The third possibility would be that non-emotional processes related to performance in instrumental conditioning are altered in ASD. ASD patients show different patterns of decision making compare to normal subjects. For example, ASD patients showed risk-aversive attitudes during the Iowa gambling task (South et al., 2014) and a computerized financial decision-making task (Gosling & Moutier, 2018). These altered decision-making processes may delay reversal learning in an instrumental conditioning paradigm. Clearly, additional studies are required to resolve these alternative possibilities.

To summarize, our results demonstrate that one consequence of *Shank2* knock-out in mice is abnormally heightened negative emotional processing that limits behavioral flexibility under certain circumstances. Our findings suggest that behavioral flexibility may be seriously limited by abnormal emotional processing in ASD. Note that behavioral inflexibility can be caused by abnormality in many different underlying processes such as inhibitory control, value-based decision making under free-choice conditions, and capturing complex stimulus-response-outcome contingencies (task rules) under cognitively demanding situations. In this regard, our results do not argue directly against the cognitive inflexibility hypothesis for ASD (Geurts et al., 2009; Van Eylen et al., 2011). Further studies are needed to determine the extent to which ASD patients and animal models show impairments in flexibly adjusting behavior under diverse behavioral settings and its underlying neural processes.

## Materials and Methods

### Subjects

We used *Shank2-*KO mice with deletions in exons 6 and 7 of *Shank2* gene which mimic the microdeletion of exons 6 and 7 in human *SHANK2* gene identified in ASD (Won et al., 2012). We used 32 male *Shank2-*KO mice and 30 male WT littermates (3~6 months old) in this study. Eighteen (nine WT and nine *Shank2-*KO) mice were used in Experiment 1, three (three *Shank2-*KO) mice were used in Experiment 2, 20 (10 WT and 10 *Shank2-* KO) mice were used in Experiment 3, and seven mice (two WT and five *Shank2-*KO) mice were used in Experiment 4. One WT mouse was used in both Experiment 1 and Experiment 4, one *Shank2-*KO mouse in both Experiment 1 and Experiment 2, and 12 (eight WT and four *Shank2-*KO) mice in both Experiment 2 and Experiment 4. Those mice used in Experiment 2 as well as another experiment were first tested in Experiment 2. Mice were bred with C57BL/6N background and characterized by PCR genotyping as previously reported (Chung et al., 2019; Won et al., 2012). For classical conditioning, mice were water-deprived and their body weights were maintained above 80% of the initial body weights after handling. Those mice used to measure eye closure responses to air puff were fed *ad libitum*. All animal care and experimental procedures were performed in accordance with protocols approved by the directives of the Animal Care and Use Committee of Korea Advanced Institute of Science and Technology (approval number KA2018-08).

### Surgery

A customized aluminum head plate was implanted on the skull under isoflurane (1.5–2.0 % in 100 % [v/v] oxygen) anesthesia. The head plate was placed near the lambdoid suture and fixed by screws and dental cement. The mice were allowed to recover > 1 week before behavioral training began.

### Classical conditioning

All mice were trained in a probabilistic classical conditioning task under head fixation (**Figure 1a**) as previously described (Jeong et al., 2020). The animal’s head was fixed to a custom-built metal holder. A water port (a blunt 17-gauge needle) was placed slightly below the animal’s nose, an air puff port (a blunt 18-gauge needle) was placed 3~5 mm away from the animal’s left eye, and an odor port (silicon tube; diameter, 8 mm) was placed slightly above the animal’s nose. Four different odors (citrus, isononyl acetate, L-carvone, and 1-butanol) were dissolved in mineral oil (1:1000, v:v) and delivered to the animal using an air circulation system. Two odors were selected randomly from the four for each experiment and for each animal. The animal’s licking behavior was detected by an infrared light sensor placed adjacent to the water port.

Behavioral phases consisted of habituation, acquisition, and reversal. For habituation, a small amount of water (6 μl) was provided from the water port initially in every 5 s without an odor cue (~50 trials, day 1). Then the same amount of water was provided 1 s after the delivery of an odor cue (1 s) that was different from those used in the acquisition phase, and a 2.5~4.5 s inter-trial interval (uniform random distribution) was imposed. The habituation phase lasted 1~3 sessions (~400 trials per session).

In the acquisition phase, an odor cue randomly chosen from two different odors was delivered for 1 s and, after a delay of 1 s, an associated outcome was delivered with a given probability (**Figure 1b**). An inter-trial interval (2.5~4.5 s, uniform random distribution) was then imposed before the next trial began. The mice were trained for three daily sessions of 400 trials in the acquisition phase. We used three different sets of cue-outcome contingency. Reward-predicting conditioned stimulus (CS_Rw_) and punishment-predicting conditioned stimulus (CS_Pn_) were paired with water (6 μl) and air puff, respectively, as the following:

Experiment 1: CS_Rw_ (75% delivery of water) and CS_Pn_ (75 % delivery of strong air puff; 100 ms, 3 psi).
Experiment 3: CS_Rw_ (75 % delivery of water) and CS_Pn_ (75 % delivery of mild air puff; 5 ms, 3 psi).
Experiment 4: CS_80%_ (80% delivery of water) and CS_20%_ (20% delivery of water).

In the reversal phase, the cue-outcome contingency of the acquisition phase was reversed. In Experiments 1 and 2, a reward-predicting cue before reversal was paired with a punishment after reversal (CS_Rw→Pn_) and a punish-predicting cue before reversal was paired with a reward after reversal (CS_Pn→Rw_). In Experiment 3, the cue predicting 80% reward delivery before reversal was paired with 20% reward delivery after reversal (CS_80%→20%_) and the cue predicting 20% reward delivery before reversal was paired with 80% reward delivery after reversal (CS_20%→80%_). Each mouse was trained until reversal criterion over 1~5 daily sessions (400 trials per session). The reversal criterion was determined as the following: the anticipatory lick rate during the delay period was smoothed with a moving window of 25 trials. The reversal criterion was the first trial since cue-outcome contingency reversal at which the smoothed anticipatory lick rate was significantly higher (*t*-test, *p* < 0.05) following CS_Pn→Rw_ than CS_Rw→Pn_ (or CS_20%→80%_ than CS_80%→20%_) presentation and remained that way for five consecutive trials.

### Eye closure response

In Experiment 2, only air puffs were delivered without odor cues, and the area of the left pupil was measured with an infra-red camera at 30 Hz as previously described (Heiney, Wohl, Chettih, Ruffolo, & Medina, 2014; Jeong et al., 2020). The measured pupil area was normalized between 0 (fully open eye) to 1 (fully closed eye). Mice were habituated to the head-fixed experimental setting for 1 h for 3 days. They then received different durations (5, 10, 50, 100 ms) and intensities (3, 7, 15, 30 psi) of air puff (total 16 combinations), five times each. Consecutive air puffs were separated by 9~11 s intervals (uniform random distribution).

### Statistical tests

Sample sizes were determined based on previous behavioral studies on ASD model mice (e.g., Won et al., 2012). All statistical tests were performed with MATLAB (version R2017a) and SPSS (version 25.0). Group comparisons were performed with Student’s *t*-tests and two-way mixed ANOVA followed by post-hoc Bonferroni tests. All statistical tests were two-tailed. Statistical significance was accepted if *p* < 0.05. All data are expressed as mean ± SEM. Raw data and code to reproduce this work are archived at Dryad (https://doi.org/10.5061/dryad.crjdfn344).

## Author Contributions

M.Y. and M.W.J. conceived the study. M.Y. collected and analyzed the data. M.W.J. and E.K. supervised the study. M.Y. and M.W.J. wrote the manuscript with inputs from E.K.

## Acknowledgements

This work was supported by the Institute for Basic Science (IBS-R002-D1 to E.K. and IBS-R002-A1 to M.W.J.).

## Figure Supplement

**Figure 3-supplement figure 1.**
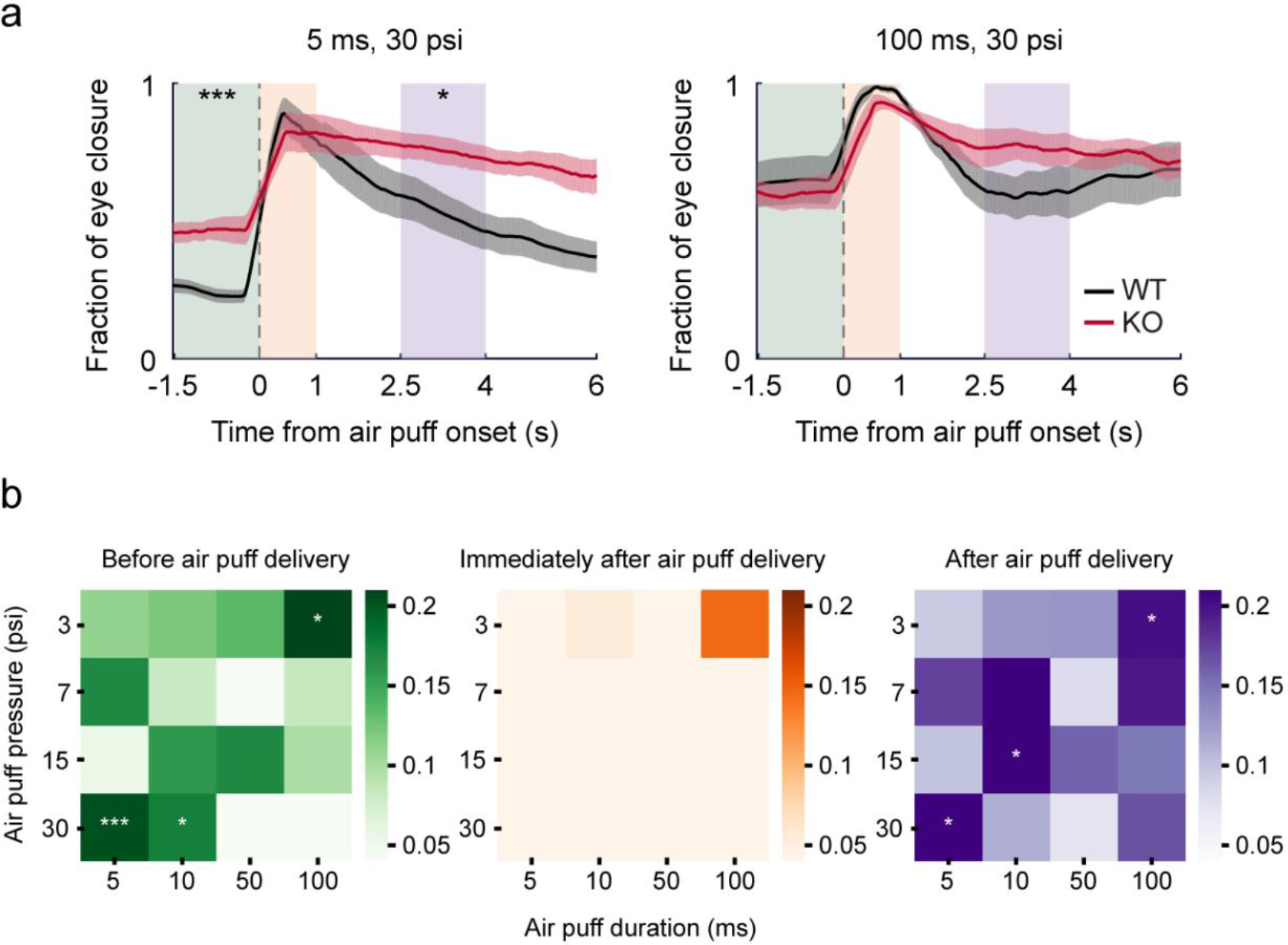
Eyelid closure responses of WT and *Shank2-*KO mice to diverse combinations of air puff pressure and duration. (**a**) Eyelid closure responses of WT (black) and *Shank2-*KO (red) mice to strong (5 ms, 30 psi) and powerful (100 ms, 30 psi) air puffs. The same format as in **Figure 3b**. (**b**) The difference in eyelid closure response between WT and *Shank2-*KO mice to 16 different combinations of air puff duration (abscissa) and pressure (ordinate) before (left; 1.5-s time window before air puff onset), immediately after (middle; 1-s time window since air puff onset), and after (right; 2.5~4 s since air puff onset) air puff delivery. Eye closure responses shown in **Figure 3** are included in these plots. \**p* < 0.05, \*\*\**p* < 0.001, *t*-test between WT and *Shank2-*KO mice (n = 8 each). Statistical test results are summarized in **Figure 3-supplement table 1**.

**Figure 3-supplement table1.** Statistical test results for eyelid closure responses to diverse combinations of air puff duration and pressure. Bold indicates a significant difference between WT and *Shank2-*KO mice (n = 8 each).

| Time window | Task Condition |  |  |  |  |
| --- | --- | --- | --- | --- | --- |
|  | Duration<br>Pressure | 5 ms | 10 ms | 50 ms | 100 ms |
| 1.5~0 s<br>before air<br>puff onset | 3 psi | $t_{(14)} = 1.003$<br>$p = 0.333$ | $t_{(14)} = 1.060$<br>$p = 0.307$ | $t_{(14)} = 1.240$<br>$p = 0.235$ | **$t_{(14)} = 2.380$**<br>**$p = 0.032$** |
| | 7 psi | $t_{(14)} = 1.608$<br>$p = 0.130$ | $t_{(14)} = 0.596$<br>$p = 0.560$ | $t_{(14)} = 0.172$<br>$p = 0.866$ | $t_{(14)} = 0.698$<br>$p = 0.497$ |
| | 15 psi | $t_{(14)} = 0.478$<br>$p = 0.640$ | $t_{(14)} = 1.872$<br>$p = 0.082$ | $t_{(14)} = 1.889$<br>$p = 0.080$ | $t_{(14)} = 0.912$<br>$p = 0.377$ |
| | 30 psi | **$t_{(14)} = 4.723$**<br>**$p = 3.0 \times 10^{-4}$** | **$t_{(14)} = 2.280$**<br>**$p = 0.039$** | $t_{(14)} = 0.203$<br>$p = 0.842$ | $t_{(14)} = 0.505$<br>$p = 0.622$ |
| 0~1 s after<br>air puff onset | 3 psi | $t_{(14)} = 0.383$<br>$p = 0.707$ | $t_{(14)} = 1.143$<br>$p = 0.272$ | $t_{(14)} = 0.713$<br>$p = 0.488$ | $t_{(14)} = 1.484$<br>$p = 0.160$ |
| | 7 psi | $t_{(14)} = 0.295$<br>$p = 0.773$ | $t_{(14)} = 0.402$<br>$p = 0.694$ | $t_{(14)} = 2.138$<br>$p = 0.051$ | $t_{(14)} = 1.390$<br>$p = 0.186$ |
| | 15 psi | $t_{(14)} = 0.741$<br>$p = 0.471$ | $t_{(14)} = 0.111$<br>$p = 0.913$ | $t_{(14)} = 1.409$<br>$p = 0.181$ | $t_{(14)} = 1.894$<br>$p = 0.079$ |
| | 30 psi | $t_{(14)} = 0.325$<br>$p = 0.750$ | $t_{(14)} = 0.271$<br>$p = 0.790$ | $t_{(14)} = 1.335$<br>$p = 0.203$ | $t_{(14)} = 1.816$<br>$p = 0.091$ |
| 2.5~4 s after<br>air puff onset | 3 psi | $t_{(14)} = 0.889$<br>$p = 0.389$ | $t_{(14)} = 1.408$<br>$p = 0.181$ | $t_{(14)} = 1.564$<br>$p = 0.140$ | **$t_{(14)} = 2.157$**<br>**$p = 0.049$** |
| | 7 psi | $t_{(14)} = 1.617$<br>$p = 0.128$ | $t_{(14)} = 2.123$<br>$p = 0.052$ | $t_{(14)} = 0.662$<br>$p = 0.519$ | $t_{(14)} = 1.865$<br>$p = 0.083$ |
| | 15 psi | $t_{(14)} = 0.913$<br>$p = 0.377$ | **$t_{(14)} = 2.305$**<br>**$p = 0.037$** | $t_{(14)} = 1.746$<br>$p = 0.103$ | $t_{(14)} = 1.419$<br>$p = 0.178$ |
| | 30 psi | **$t_{(14)} = 2.598$**<br>**$p = 0.021$** | $t_{(14)} = 1.173$<br>$p = 0.261$ | $t_{(14)} = 0.811$<br>$p = 0.431$ | $t_{(14)} = 1.717$<br>$p = 0.108$ |

